# Pardon the interruption: saccadic inhibition enables the rapid deployment of alternate oculomotor plans

**DOI:** 10.1101/285841

**Authors:** Emilio Salinas, Terrence R. Stanford

## Abstract

Diverse psychophysical and neurophysiological results show that oculomotor networks are continuously active, such that plans for making the next eye movement are always ongoing. So, when new visual information arrives unexpectedly, how are those plans affected? At what point can the new information start guiding an eye movement, and how? Here, based on modeling and simulation results, we make two observations that are relevant to these questions. First, we note that many experiments, including those investigating the phenomenon known as “saccadic inhibition,” are consistent with the idea that sudden-onset stimuli briefly interrupt the gradual rise in neural activity associated with the preparation of an impending saccade. And second, we show that this stimulus-driven interruption is functionally adaptive, but only if perception is fast. In that case, putting on hold an ongoing saccade plan toward location *A* allows the oculomotor system to initiate a concurrent, alternative plan toward location *B* (where a stimulus just appeared), deliberate (briefly) on the priority of each target, and determine which plan should continue. Based on physiological data, we estimate that the actual advantage of this strategy, relative to one in which any plan once initiated must be completed, is of several tens of milliseconds.

---

A fundamental function of oculomotor circuits is to determine where the eyes should look next and produce the appropriate eye movement. Neurally, each saccade is the culmination of a motor planning process whereby the firing rates of movement-related neurons rise gradually, monotonically, until the ramping activity of the population reaches a particular threshold level, at which point the plan becomes an uncancelable command and the saccade is triggered^1,2,3,4,5,6^. A timing conflict is likely to arise because this rise-to-threshold process unfolds over a sizable period of time, but new, potentially critical visual information may arrive at any moment. Furthermore, due to two prominent features of oculomotor circuits, this conflict must be extremely common.

On one hand, both behavioral and neurophysiological evidence indicates that saccades are programmed continuously, which is to say that there is always a subset of movement-related neurons (encoding a particular movement vector) that is steadily increasing its activity toward threshold. First, both humans and monkeys generate saccades at similar rates under a wide variety of viewing conditions^7,8,9^, even in the dark^10,11^. Second, activity recorded in the frontal eye field (**FEF**) during free-viewing conditions shows, from one saccade to the next, variations in firing rate occurring around a relatively high mean compared to that seen after prolonged fixation^9,12,13^, as if the movement-related activity never fell far below threshold. Third, motor plans are not contingent on the completion of a target selection process^5,14,15,16,17^ and may proceed in parallel^5,6,9,16,18,19,20^. And fourth, the streaming of motor plans is strongly driven by current task contingencies and behavioral goals^21,22^; that is, by internal, already acquired information, and not necessarily by the visual stimuli currently in view.

On the other hand, saccades are rapidly and preferentially drawn to physically salient or highly relevant visual stimuli^8,23,24,25,26,27,28,29,30^, such as a traffic light turning red, or a mouse suddenly scurrying across the kitchen floor. The draw of such stimuli is often described as “attentional capture” (refs. 31, 32) or, when particularly potent, “oculomotor capture” (refs. 25, 26) Stimuli that capture attention engage dedicated neural mechanisms that are highly sensitive to bottom-up information and are closely related to the selective firing of visually-responsive neurons within oculomotor circuits^20,33,34,35,36,37^.

Therefore, it must be typically the case that, when a salient stimulus appears abruptly, the corresponding visual information arrives at oculomotor planning areas while the activity associated with some movement vector is already rising toward threshold. What should the system do then? Should the ongoing plan be completed first, before the new information is acted upon, or should the plan be canceled and replaced with another one toward the recent stimulus? Clearly, the answer depends on the priorities of the targets. So, it may be best to put the ongoing saccade plan on hold, evaluate the new sensory information, and *then* determine whether to continue with the current plan or to cancel it in favor of an alternate one.

This intuition has been mentioned earlier^38,39,40^. For instance, Anderson and colleagues^40^ articulated the problem this way: “involuntary attentional capture can rapidly orient the organism to unexpected changes that could signal danger or opportunity, but has the potential to cause distraction from intended acts of perception.” That is, delaying a response to a visual stimulus that appears abruptly could be costly when it reveals an imminent threat or a fleeting opportunity, but always canceling the ongoing saccade plans would be quite inefficient, because not all salient stimuli have such importance. Here, we provide an initial framework for evaluating this intuitive tradeoff quantitatively. First we show that a simple descriptive model replicates a variety of experimental results which, in effect, indicate that ongoing saccade plans are transiently interrupted by salient, sudden visual stimuli. Then, based on this model, we use analytical and numerical calculations to determine the conditions under which such interruption is behaviorally advantageous, and estimate the actual temporal tradeoff that it entails.

## Results

### Behavioral manifestation of an interruption in motor planning

First we make a simple observation, which is that, across multiple trials, a transient interruption in the rising activity that comprises the motor plan to make a saccade leaves a characteristic signature in the corresponding distribution of saccadic RTs — a dip.

The neural events that precede the onset of a saccade are well understood, particularly when the eye movement is in response to the presentation of a visual target. After the target is shown, activity in oculomotor areas, most prominently the FEF and superior colliculus (**SC**), starts building up, with the activated neurons encoding the movement vector required to fixate the target. The ramping activity keeps increasing until it reaches a point of no return — the threshold — and shortly thereafter (∼10–20 ms) the saccadic movement is initiated^1,2,3^. Although the baseline and threshold levels of this rise-to-threshold process may vary and contribute substantially to its dynamics^6^, in general, RTs are most strongly correlated with variations in the build-up rate^1,4,5,6,16,41^. Thus, a reasonable simplification is that short versus long RTs largely correspond to steeper versus shallower excursions in activity between baseline and threshold (Fig. 1a).

**Figure 1.**
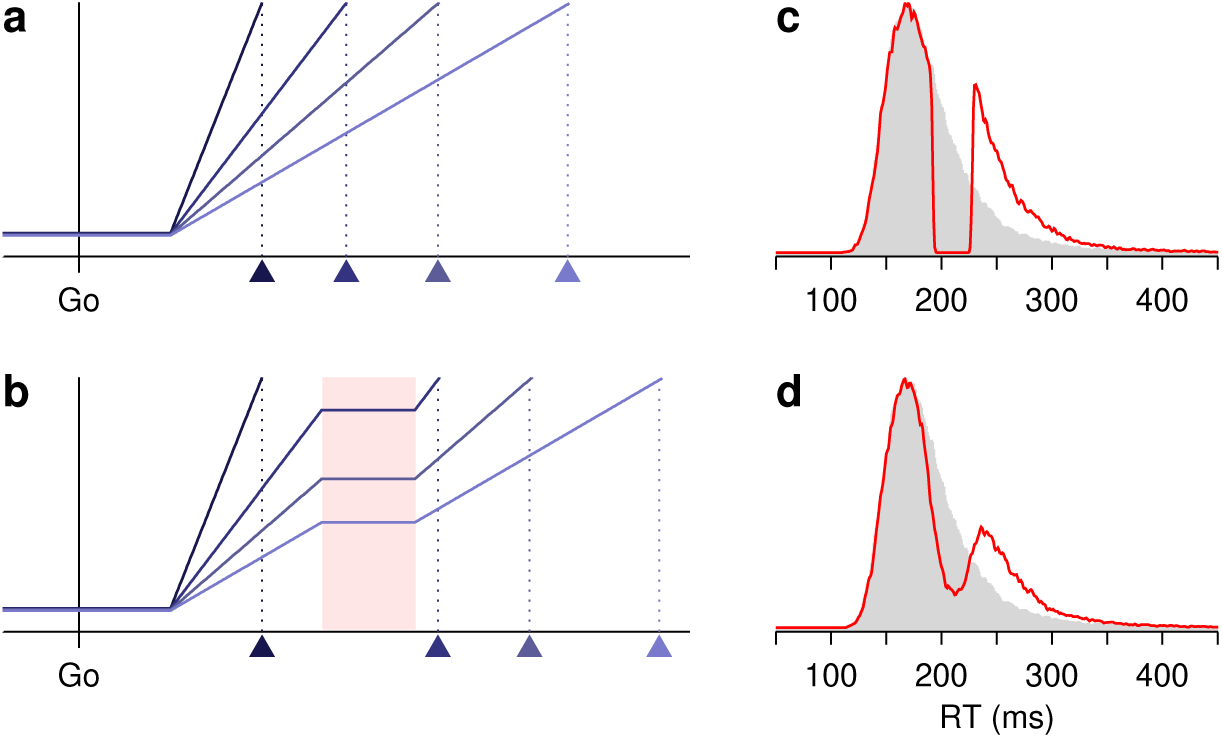
Interruptions in motor planning produce a dip in the distribution of saccadic RTs. **a**, Four examples of the rise-to-threshold process. The y axis represents oculomotor activity as a function of time. Activity increases gradually, and when a critical threshold level is reached, a saccade is triggered. All four plans are identical except for their build-up rates. Triangles indicate saccade onset. **b**, Four example motor plans that are briefly interrupted. The rise in activity halts during the interruption period (red shade). Build-up rates are the same as in the panel above. **c**, Simulated RT distributions for motor plans that rise to threshold uninterrupted (gray shade) or that halt (red line) for 36 ms (between 192 and 228 ms after the go signal) but are otherwise identical. Distributions were obtained from 50,000 simulated trials using 1 ms RT bins and Gaussian smoothing with *σ* = 1 ms. Note the sharp discontinuity produced by the pause. **d**, As in **c**, except that the probability of interruption for any given trial was equal to 0.7 (rather than 1), and the onset and offset of the interruption interval varied normally with a SD equal to 8 ms (instead of 0). Note the smooth dip in the distribution.

In agreement with this account, it is well established that the RT distributions of simple reactive saccades, which have a characteristic skew, are accurately replicated by a linear rise-to-threshold process in which the build-up rate, *r*_*B*_, is constant within each trial but is drawn from a Gaussian distribution from one trial to the next^42,43^. Under such conditions, the RT can be expressed as

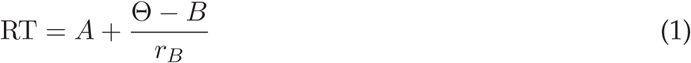

where *A, B*, and Θ are constants (the afferent delay, baseline activity, and threshold), and for each trial the build-up rate, *r*_*B*_, is a different sample from the same Gaussian distribution. Simulated RT distributions based on this expression (Fig. 1c, d, gray shades) closely approximate distributions obtained experimentally based on saccades to single targets.

This is the basis for our model, which is simply a descriptive one. It provides a minimalistic depiction of the underlying neural activity, and is entirely agnostic regarding the underlying circuit dynamics and mechanisms (in contrast to, for instance, the models considered by Bompas and Sumner^44^). Also, it is similar in many ways to other models that make slightly different assumptions (e.g., refs. 43, 45). Importantly, however, the results do not depend on these minor differences. The key property — in agreement with the underlying neurophysiology — is that the variance in RT is largely due to the variance in build-up rate.

What we wish to point out is the characteristic effect produced when the rise-to-threshold process is interrupted during a consistent time period defined with respect to the onset of the rise. If the build up of activity is momentarily halted, such that the firing level is maintained constant during the pause (Fig. 1b), then the RT distribution is essentially split into two pieces separated by an empty interval (Fig. 1c, red trace). The location of the void within the distribution depends on the temporal offset between the go signal (or more precisely, the start of the rise to threshold) and the onset of the interruption, and the length of the void is equal to the duration of the pause. Whatever the offset, if the two parts of the distribution were brought together, the original distribution without interruption would be recovered. Thus, ideally, the behavioral manifestation of a consistent interruption of the saccadic motor plans is a discontinuous split of the RT distribution.

Notably, though, any variability in the interruption mechanism will turn the discontinuous split into a smooth dip (Fig. 1d). (The sharpness of the discontinuity will also depend on the bin width used to generate the RT histograms, but this effect should be minor.) In particular, there are three ways in which such smoothing would be likely to occur. (1) Rather than halting, the motor plan could keep increasing at a low build-up rate, i.e., it could be just partially suppressed. A nonzero build-up rate would partially fill in the void in the RT distribution, and if that low rate fluctuated randomly across trials, the edges of the void would be softened. (2) Rather than always halting, the motor plan could halt on some trials but not on others. That is, the probability of halting could be less than 1. (3) Rather than being constant, the duration of the interruption (and/or its onset and offset) could also fluctuate randomly across trials. As with point 1, the larger the fluctuations, the heavier the smoothing. In combination, these factors can easily turn a sharp, discontinuous split in the RT distribution into a visible but much more subtle, smooth dip (compare red traces in Fig. 1c, d).

The effect of the interruption can also be appreciated based on how much each individual RT is delayed. If on a given trial the interruption halts the motor plan starting at a time *I*_*on*_ and lasts *q* ms, then the observed RT is simply

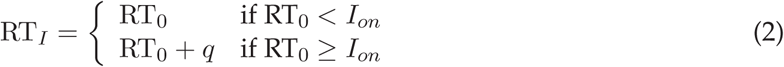

where RT_0_ is the value that would have been measured had the same saccade plan (with the same build-up rate) not been interrupted. In other words, as can be seen in Fig. 1b, if the interruption starts at *I*_*on*_ (left edge of red shade), saccades that are programmed to occur before *I*_*on*_ are executed normally, whereas saccades that are programmed to occur after *I*_*on*_ are delayed by *q* ms, which is the duration of the pause. This is true on a trial-by-trial basis. So, assuming that the activity level does not change during the interruption and that *I*_*on*_ and *q* are constant, equation (2) leads to a sharp break in the RT distribution that starts at *I*_*on*_ and is *q* ms wide. In contrast, when *I*_*on*_ and *q* are not constant, the fluctuations across trials simply smooth the edges of the otherwise discontinuous, empty interval. Notably, because the above expression applies to any RT_0_, the break or dip occurs regardless of the shape of the original RT distribution.

### Saccadic inhibition as an interruption

Many psychophysical studies have characterized the phenomenon best known as “saccadic inhibition” or the “remote distracter effect,” in which the sudden presentation of a visual stimulus delays the execution of a saccade. In this section we show that, although the results of these studies are typically not described in terms of an interruption in motor planning, that is precisely what they reveal.

Saccadic inhibition is typically produced by presenting a brief distracting stimulus while a participant is just about to make a saccade to a target (for example, Fig. 2a). Over multiple trials, the influence of the distracter manifests as a distribution of saccadic RTs (for correct responses) with a dip that depends on the timing of the distracter (for example, Fig. 2d–g, black traces) — a signature of a pause in the rise-to-threshold process that comprises a saccadic motor plan, as discussed above. As evaluated by this signature, saccadic inhibition is extremely robust.

**Figure 2.**
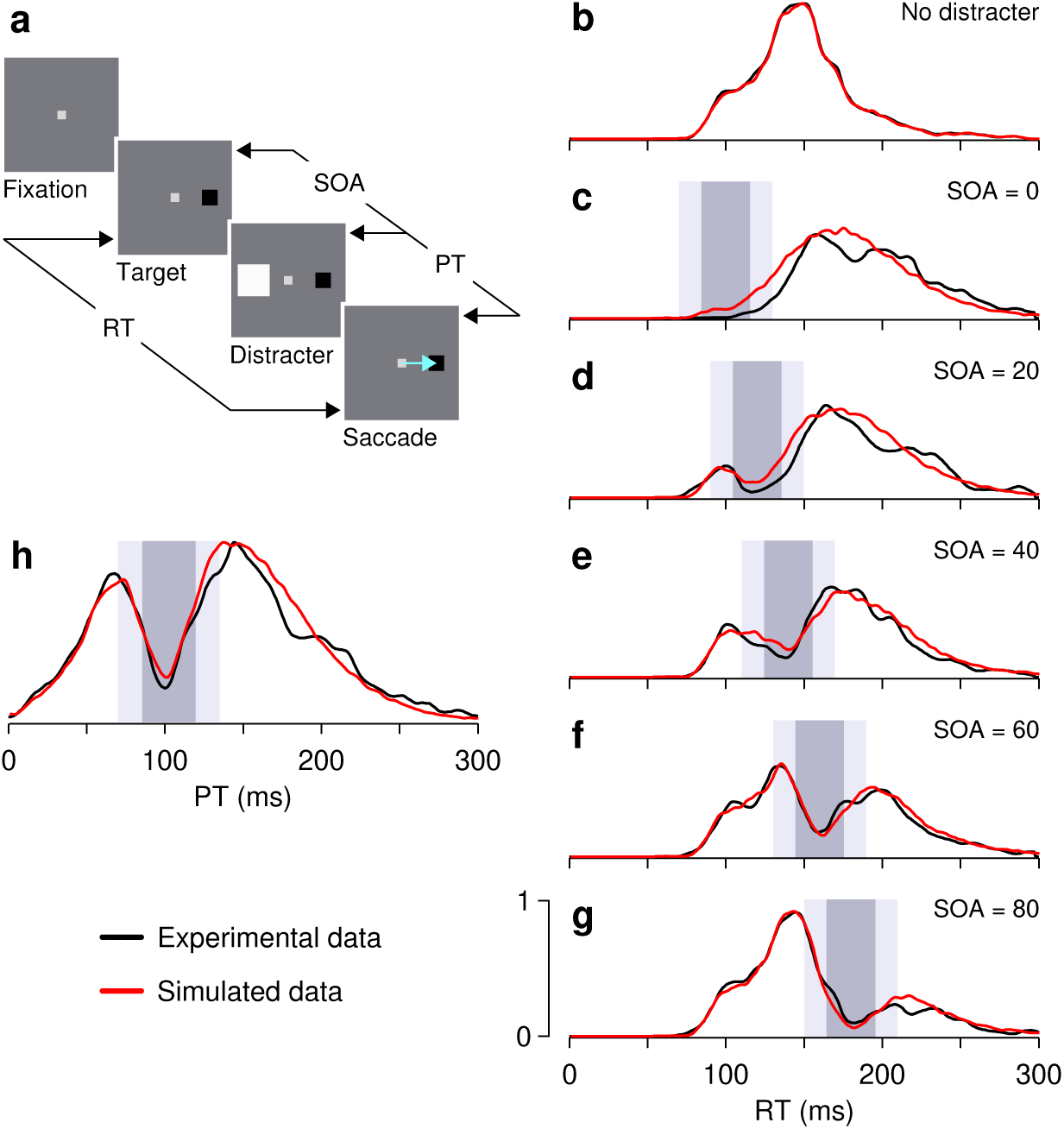
Interruptions in motor planning observed experimentally. **a**, The paradigm used by Bompas and Sumner^44^ to study saccadic inhibition. Participants were instructed to make a saccade (blue arrow) to the target (black square) as soon as it appeared, and to ignore any distracters. In most trials, a distracter (white square) was briefly flashed (50 ms) opposite to the target. The delay between target onset and distracter onset (SOA) varied across trials. For each trial, the PT is equal to RT - SOA, and corresponds to the time interval between distracter onset and saccade onset. **b**–**g**, RT distributions reported by Bompas and Sumner^44^ (observer 1, black traces) along with our model results (red traces). SOAs are indicated, in ms. In the rise-to-threshold model, RT_0_ samples were directly drawn from the distribution obtained in the no-distracter condition (b). The effect of the interruption was then calculated for each drawn RT_0_ using equation (2), taking the corresponding SOA into account. The interruptions occurred, on average, between 85 and 115 ms after distracter onset (dark gray shades) with a probability of 1, but the onset and offset times varied across trials (SD was 14.3 ms, correlation was −0.8; light gray shades show 1 SD in each direction). **h**, PT distributions for the simulated (red trace) and experimental data from Bompas and Sumner^44^ (observer 1, black traces). PT histograms include aggregated data from all SOAs. All experimental data were redrawn from Bompas and Sumner^44^.

The characteristic dip is observed whether the distracter stimulus is discrete and localizable (e.g., refs. 39, 44) or widely distributed and lacking a well-defined spatial location (e.g., wide bars flashing above and below the intended saccade target^38^). The strength of the effect depends on the saliency of the distracting event; for instance, it is stronger for larger^46,47,48^ and higher contrast stimuli^44^. Variations in other stimulus dimensions, such as spatial frequency, produce only small variations (∼10 ms) in the leading phase of the dip^49^. And notably, the effect is stronger and more prolonged when the stimulus appears in an attended or task-relevant location as opposed to an unattended, task-irrelevant one^46,50^. Thus, all manner of abrupt visual stimuli produce saccadic inhibition, but their saliency matters.

In addition, the phenomenon is surprisingly independent of volitional control and of the way in which saccades are triggered. For instance, similar effects on RT are produced whether participants are instructed to make a saccade to the target or an antisaccade away from it^38^. Consistent with this, robust saccadic inhibition is observed in the context of more cognitively demanding behaviors as well, such as search tasks^49^, double-step saccades^39^, and reading^46^. At the other extreme, saccadic inhibition is equally robust for reflexive saccades observed during the quick phase of nystagmus, both during its normal operation^51^ (optokinetic nystagmus) and when it is pathological^52^ (infantile nystagmus). How the saccade plan is initiated does not seem to matter; the interruption cannot be prevented.

The robustness of the phenomenon can be appreciated based on representative RT data from an experiment by Bompas and Sumner^44^ in which human participants made saccades to single targets and were explicitly instructed to ignore any distracting stimuli, which were briefly flashed at a location diametrically opposite to the target (Fig. 2a). In trials with no distracter, the distribution of saccadic RTs was unimodal, with the expected long tail on the right side (Fig. 2b). In trials in which a distracter was shown, the resulting RT distributions demonstrated a clear dip, with the minimum consistently occurring about 100 ms after distracter onset. As the stimulus onset asynchrony (**SOA**) between target and distracter increased, the dip in the RT distribution shifted further to the right, exactly as expected from an interruption in motor planning that is time-locked to the onset of the distracter (Fig. 2c–g, black traces). Interestingly, for an SOA of 0 ms, the effect looks less like a split and more like a pure rightward shift of the whole distribution –which is exactly what one would expect from an interruption that occurs just at the onset of the rise-to-threshold process. The resulting progression goes from the original, unimodal RT distribution without any distracters, to the shifted distribution at the shortest SOA, to distributions with a pronounced dip that gradually shifts to the right.

This progression was accurately reproduced by a simple rise-to-threshold model (see Methods) in which the parameters of the interruption are constant and the only element that changes is the onset of the interruption, as dictated by the SOA (Fig. 2b–g, red traces; see caption). This simulation made no assumptions about the underlying distribution of build-up rates; it simply drew samples from the actual experimental distribution of RTs measured by Bompas and Sumner^44^ in the no-distracter condition (Fig. 2b), and for each sample calculated the effect of the interruption using equation (2). Thus, both the statistics of the build-up rate and the statistics of the interruption used to generate the synthetic data were always the same (Fig. 2, caption). The simulated interruption produced dips at different points of the RT distribution depending only on the SOA –precisely as observed in the experimental data.

Having set the interruption parameters to fit the RT distributions (Fig. 2c–g), the simulated data were then aligned to the onset of the distracter and combined across SOAs to generate a single response histogram. For each simulated trial, we calculated the interval between distracter onset and saccade onset, which we refer to as the processing time, or **PT** (Fig. 2a). This is the maximum amount of time that the system has to process the distracter signal in each trial. The resulting PT distribution (Fig. 2h, red trace) revealed a more prominent dip, and the dip observed in the experimental distribution closely matched the expectation (Fig. 2h, black trace). The fact that the experimental PT histogram, which combines data across all SOAs, indeed reveals a pronounced dip is further evidence that the interruption is synchronized to the presentation of the distracter. Curves like those in Fig. 2h are reported in many studies^38,39,46,47,48,49,50,51,52^. The point here is simply that the transient interruption in motor planning provides a compact and accurate phenomenological description of the empirical data.

Some direct neurophysiological evidence is also available indicating that a sudden-onset distracter briefly inhibits the progress of an ongoing saccade plan. In the SC, presenting an irrelevant distracter while a monkey is preparing to make an eye movement to a known location produces a brief decrease in the activity of visuomotor neurons precisely at the time expected based on the behavioral studies, about 90 ms after distracter onset^53,54^.

In conclusion, plentiful evidence indicates that the oculomotor system responds reflexively and rapidly (within ∼60 ms) to sudden changes in visual input in a manner that is consistent with ongoing saccade plans making a brief pause in their rise toward threshold.

### Temporal advantage of saccadic inhibition

Under what conditions is it advantageous to halt an ongoing saccade plan? And what is the actual temporal advantage? We answer these questions by comparing different strategies for updating and prioritizing saccade plans toward two targets.

Suppose that a stimulus appears at location *B* while a saccade plan toward location *A* is already ongoing. Then, if the stimulus is indeed relevant, i.e., worth examining right away, how quickly can a saccade to it be generated? We consider two simplified scenarios, one in which saccades cannot be interrupted and another in which they can (Fig. 3).

**Figure 3.**
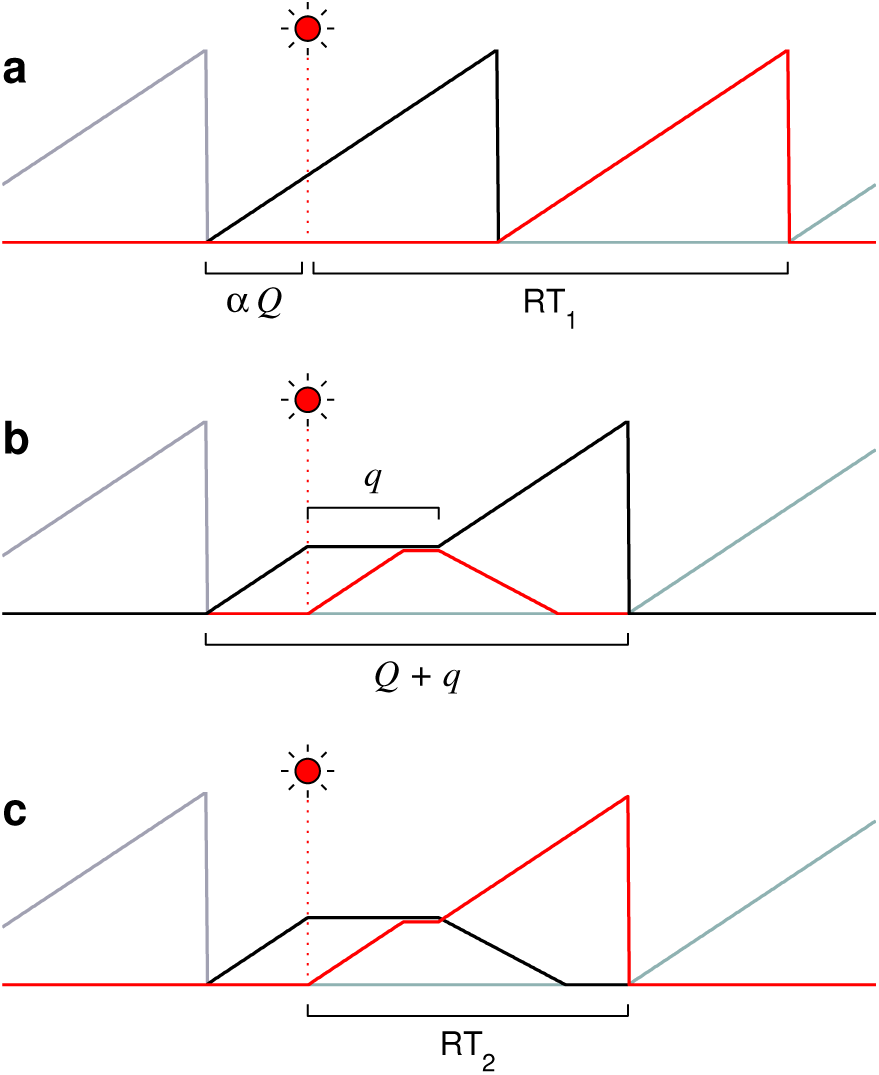
Two scenarios whereby saccade plans may incorporate new visual information. Traces represent the activity of saccade-related neurons, with colors corresponding to populations encoding distinct saccade vectors. Activity rises progressively, and when it reaches threshold, a saccade is triggered and a new motor plan is initiated. Here, all plans have the same build-up rate. The circuit detects a new stimulus (red flash) at a time *αQ* relative to the onset of the ongoing plan (black trace). **a**, A sequential scenario. Saccade plans cannot be interrupted and are produced every *Q* ms. A saccade to the flash (red trace) is made after RT_1_ ms from the time of detection. **b, c**, A parallel-programming scenario. After stimulus detection, the ongoing plan (black trace) is halted for *q* ms and a second plan, toward the stimulus (red trace), starts rising. After the interruption, either the second plan is canceled (**b**), or the first one is (**c**), in which case a saccade to the flash is made after RT_2_ ms from the time of detection.

First, note that if the arriving visual signal is spatially congruent with the ongoing plan (i.e, *B* ≈ *A*), then there is no conflict and the plan should simply continue. In that case canceling the current plan and starting over would be unnecessary and waste time. And indeed, experimental evidence^47,55,56^ indicates that saccadic inhibition demonstrates the expected dependence on *A* and *B* (see Discussion). Here we consider the problem that arises when the new visual signal and the ongoing plan are spatially incongruent, in which case the best course of action will depend on the priorities of the two potential visual targets.

In the two scenarios considered, with and without interruption, movement-related activity rises toward a threshold level Θ with a build-up rate *r*_*B*_, so that the time between saccades is normally *Q* = Θ*/r*_*B*_. For simplicity, we first consider *r*_*B*_ to be constant; later we will relax this assumption (and others) and show that additional, intrinsic randomness in the build-up rate does not affect the argument or the conclusions. In the first scenario (Fig. 3a), saccades are generated every *Q* ms, one after another. When a new visual stimulus is detected at location *B*, the ongoing plan toward location *A* (black trace) needs to be completed first (generating saccade 1) before a saccade to the stimulus can be programmed (saccade 2; red trace). Assuming that the visual signal is detected by the circuit at a time *αQ* from the start of the original plan and that no additional time is required to process it, the latency for making a saccade to the stimulus at location *B* is

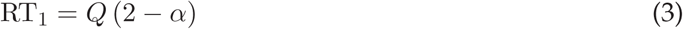

counting from the moment of detection. Note that *α* varies between 0 and 1, and indicates when the stimulus is detected by the circuit relative to how advanced the ongoing motor plan is.

In the second scenario (Fig. 3b, c), the movement-related activity also rises with a build-up rate *r*_*B*_, so in the absence of new visual information saccades are still produced every *Q* ms. However, when a new visual stimulus is detected at location *B*, again at time *αQ*, two things happen: first, the ongoing plan toward *A* is halted for *q* ms, and second, a new, competing plan toward the stimulus starts rising immediately, with the same build-up rate, *r*_*B*_. Importantly, during the *q* ms that the pause lasts, this second, parallel plan can only rise up to the level of the first plan, not further. Thus, the second plan can either catch up with the first one or stay below it — but cannot overtake it. Finally, after the interruption, two outcomes are possible: either the initial plan toward *A* continues and the budding plan toward *B* is canceled (Fig. 3b), or the reverse, the initial plan is canceled and the later one, toward the stimulus at location *B*, keeps advancing (Fig. 3c). In the former case, which occurs when the stimulus is deemed of low priority, the RT associated with the resulting saccade expands from *Q* to *Q* + *q* ms, but the eyes still land at the original target location, *A*. In the latter case, which occurs when the stimulus turns out to be of high priority, the latency for making a direct saccade to it, toward location *B*, is

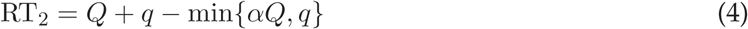

again relative to the moment of detection, where min{*a, b*} is the minimum of *a* and *b*. The question is, which scenario is more efficient?

Note that, implicitly, the time required for the stimulus to be processed in scenario 1 is conservative, in that it favors the purely sequential strategy. This is because even in the extreme case, when the stimulus is detected just before the saccade to *A* is triggered, no additional deliberation time is required. The plan toward location *B* is initiated right after the saccade to *A*, so the implicit assumption is that the perceptual evaluation of the stimulus is always completed while the first motor plan is ongoing, however short that interval may be. In contrast, in scenario 2 the interruption time, *q*, is equal to the deliberation time, i.e., the amount of time needed to resolve whether the stimulus at *B* is of low or high priority. During the pause, the plan toward *A* is put on hold and the alternate one, toward *B*, is initiated “just in case,” but the usefulness of the latter is defined only after the *q* ms have elapsed.

For a saccade to the stimulus, the difference in latency between the two scenarios is

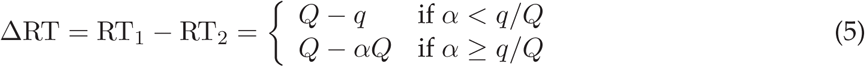

where a positive value indicates a faster response in scenario 2, in which the ongoing plan is interrupted. This expression shows that the advantage in RT depends on when the stimulus is detected relative to how advanced the ongoing plan is, which is what *α* represents. If the first plan is just starting to rise (*α* ≈ 0), the difference is potentially large; whereas if the planned saccade toward *A* is just about to be executed (*α* ≈ 1), the difference is minimal. The mean difference, averaging over detection times (i.e., integrating over *α*, assuming it is uniformly distributed), is

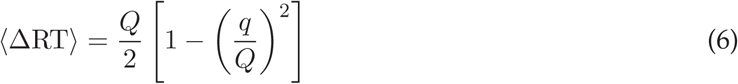

This is the main result. According to this expression, re-prioritizing on the fly pays off only when the perceptual evaluation time, *q*, is shorter than the intersaccadic interval, *Q*, in which case a positive average difference in RT is obtained. The precise advantage, however, depends on the duration of the interruption relative to the typical RT (Fig. 4a).

**Figure 4.**
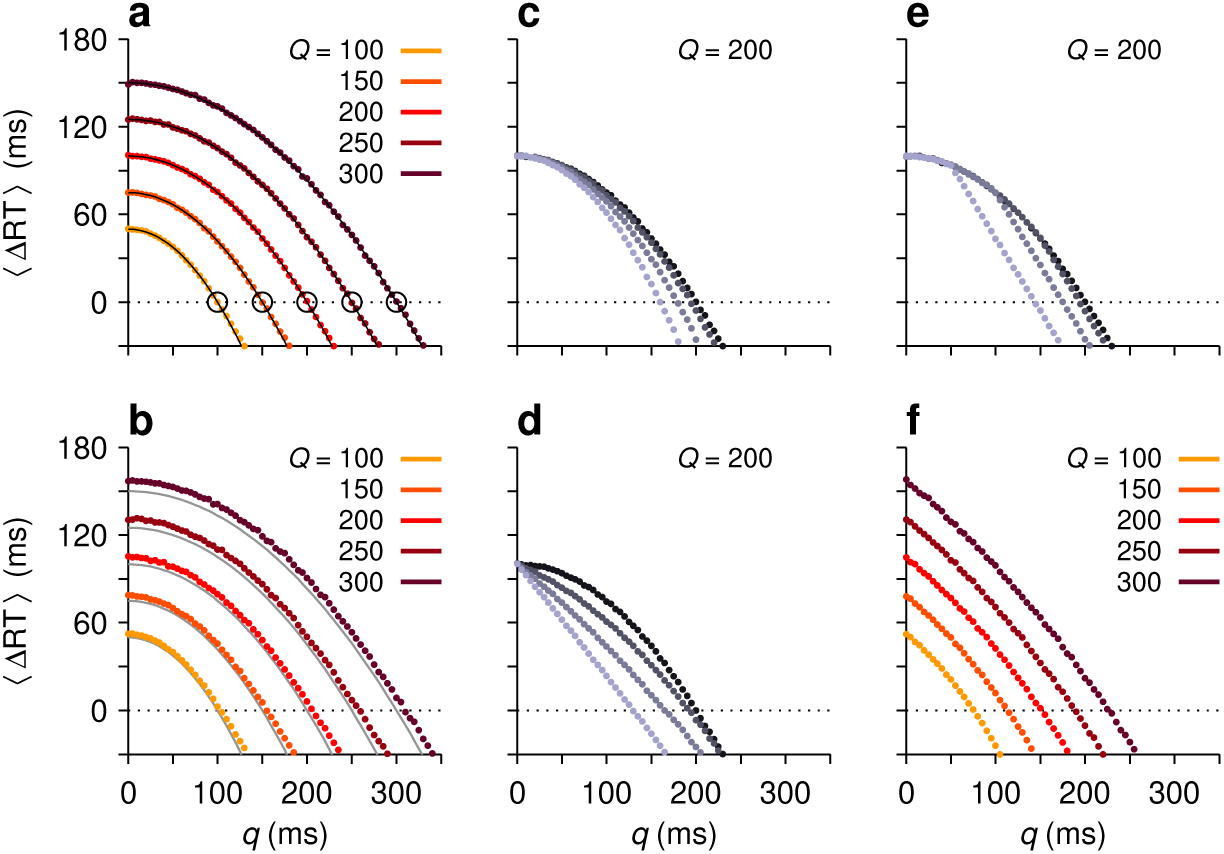
Temporal advantage of an interruption in motor planning. Each panel shows the mean difference in RT between a sequential strategy, in which plans are not interrupted, and a parallel strategy in which saccade plans are interrupted and deployed concurrently. The x axes show the deliberation time, *q*. The mean saccade latency, *Q*, is indicated. **a**, Expected difference in RT from theoretical calculations. Colored dots are results from computer simulations; gray lines are from equation (6). Circles mark the points beyond which concurrent programming ceases to be advantageous. **b**, As in **a**, but with Gaussian variability in the simulated build-up rates, such that the SD of *Q* was approximately equal to *Q/*4. **c**, Results when the ongoing motor plan is not fully arrested but keeps rising at lower build-up rate. Going from dark to light points, the build-up rate, *r*_*B*_, of the ongoing plan was equal to 0, 0.25, 0.5 and 0.75 times the initial value. **d**, Results when the concurrent motor plan toward the stimulus rises at a lower build-up rate. Going from dark to light points, the build-up rate, *r*_*B*_, of the second plan was equal to 1, 0.75, 0.5 and 0.25 times the maximum value. **e**, Results when the concurrent motor plan toward the stimulus can rise only up to a point below threshold. Going from dark to light points, the maximum rise is equal to 1, 0.75, 0.5 and 0.25 times the maximum value, Θ. **f**, As in **b**, but with all the effects in panels **c–e** with coefficients of 0.5 applied simultaneously.

What are physiologically plausible values for these two numbers? First consider an intersaccadic interval of 200 – 250 ms, which is most common in humans and monkeys^7,8,9,10^, and an interruption time of approximately 70 ms, as estimated from saccadic inhibition experiments^38,44,46,48^. Based on equation (6), those numbers give a mean difference (ΔRT) between 88 and 115 ms favoring the parallel programming strategy, which is quite large as a proportion of the median intersaccadic interval (∼45%). There is a cost, of course, which is that when the stimulus turns out to be irrelevant, the original RT is lengthened by *q* ms.

This calculation, however, might overestimate the true advantage, because saccades *can* be produced considerably more quickly (see Discussion), and because the interruption could conceivably be longer, depending on the location and saliency of the stimulus, as well as on the robustness of the motor plans. Assuming that conditions are optimized for producing short saccadic latencies (*Q* ≈ 150 ms), as happens when target locations are predictable and subjects are highly motivated^6,28,35,57^, and that the interruption is as long (*q* = 115 ms) as it could be under extreme circumstances^6,39^, the average time saved according to the above expression is now 31 ms. This is considerably smaller than above — but still sizable as a proportion of the time between fixations (∼21%).

More generally, the dependence of ⟨ΔRT⟩ on *q* and *Q* indicates that interrupting the ongoing saccade plan and initiating a parallel plan while deliberating on the priority of the new information is clearly advantageous over a wide range of physiologically relevant values (Fig. 4a) — again, as long as the perceptual processes underlying the deliberation (e.g., stimulus detection and identification) are at least as fast as the mean intersaccadic interval. Indeed, perceptual processing speeds typically satisfy this requirement, as elaborated in the Discussion.

One could worry, though, that the analytical result involves strong simplifications, so we made similar comparisons based on computer simulations in which various assumptions within the two scenarios were relaxed. As a check, we first simulated the exact conditions assumed in the above calculations, and indeed, the analytical and numerical results were in agreement (Fig. 4a, lines vs. dots). Then we allowed the build-up rate to be different for each saccade plan (*r*_*B*_ was drawn from a Gaussian distribution), adjusting the variance in build-up rate so that the corresponding saccade latency distributions were wide (i.e., so that the SD of *Q* was approximately equal to *Q/*4). Introducing a large amount of variability had a very small effect that tended to increase the advantage of the concurrent programming strategy (Fig. 4b, compare dots vs. gray lines).

Next, we relaxed three key assumptions of the second scenario in ways that tended to lessen its advantage. (1) The interruption did not fully arrest the ongoing plan, but simply diminished its build-up rate, and if the ongoing plan reached threshold before the deliberation was over, the plan toward the stimulus had to restart from the baseline. (2) The plan toward the stimulus could rise during the interruption interval but slowly, at a fraction of its nominal build-up rate. (3) The plan toward the stimulus could only rise so far during the interruption; that is, as before, the second plan could not overtake the first one, but in addition, it was not allowed to increase beyond an absolute level below the saccade threshold. All of these manipulations curtailed the amount of progress that the concurrent plan toward the stimulus could make during the pause — but in all cases the effects were gradual, not qualitative, and typically required large changes in the parameters, on the order of 50%, to be substantial (Fig. 4c–e). Even in a worst-case scenario, in which all of these alterations were introduced at the same time, the results still showed a frank advantage of the interruption mechanism over strict sequential programming within a large range of interruption and mean saccade durations (Fig. 4f).

These results show that the known physiological measurements of saccadic latencies and decision times are largely consistent with a mechanism whereby ongoing saccade plans are put on hold and re-prioritized on the fly, enabling significantly faster saccadic responses toward recently detected stimuli, with the benefit being on the order of several tens of milliseconds per saccade.

### Pronounced interruptions in the double-step task

We consider another example of an experiment in which there is strong evidence for a stimulus-driven interruption in motor planning. This case is interesting because it illustrates quite specifically how the interruption and concurrent motor programming can directly determine a subject’s performance in a well-known oculomotor paradigm, the double-step task.

We discuss the version of the double-step task implemented by Buonocore and colleagues^39^ Fig. 5a). All trials start in the same way, with the participant fixating on a central point flanked by two empty circles, or placeholders, one on the left and another on the right. Target presentation corresponds to one of the circles being filled. In one half of the trials (control trials), a target appears and the participant must simply make a saccade to it. In the other half of the trials (actual double-step trials; Fig. 5a), a target appears but then steps to the opposite location after a short delay (SOA). The participant is instructed to always look at the target; however, depending on the SOA and the participant’s urgency, the resulting eye movement may be toward the first (incorrect) or the second (correct) target location.

**Figure 5.**
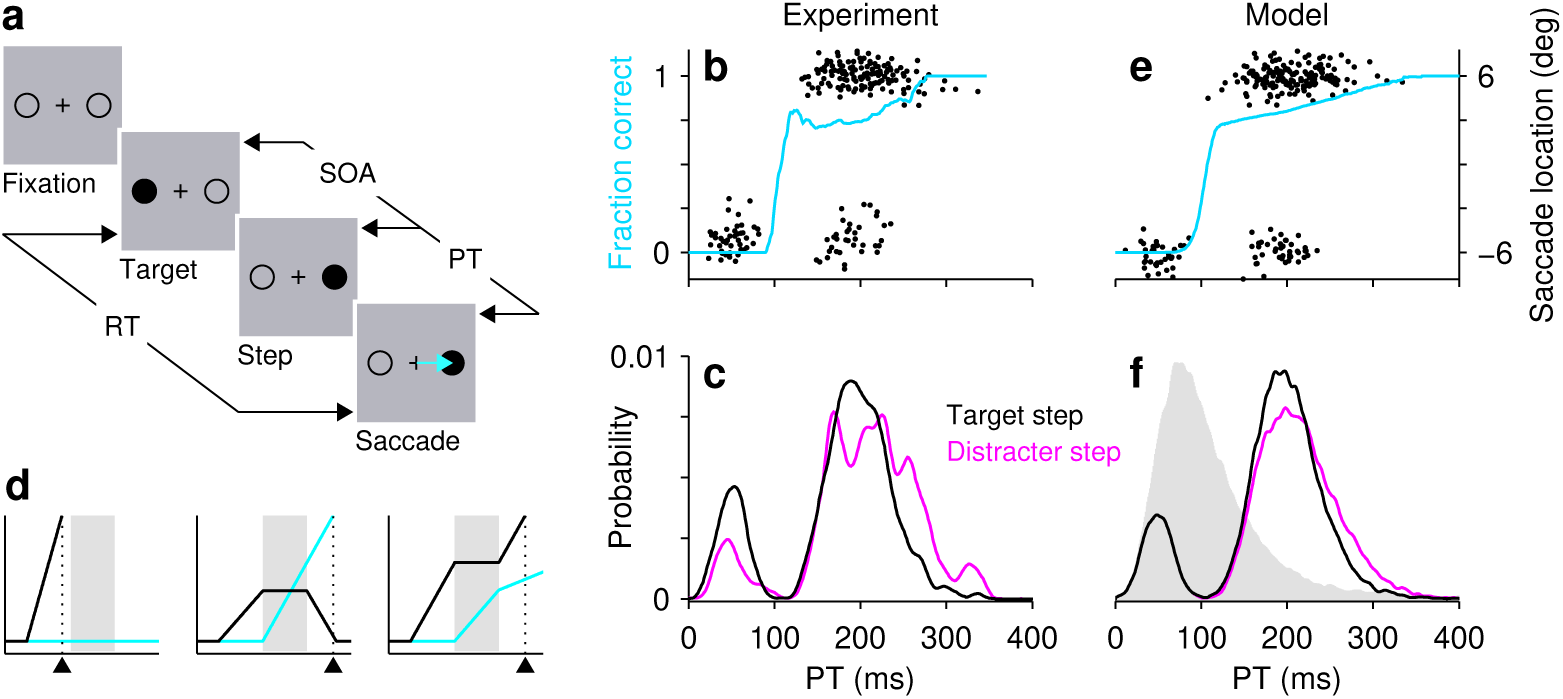
Interruptions in motor planning in the double-step task. **a**, The double-step paradigm used by Buonocore et al^39^. Participants were instructed to make a saccade (blue arrow) to the target (filled circle) as soon as it appeared. In one half of the trials the target did not change (not shown); in the other half it stepped to the opposite location after a delay (SOA). The PT in each trial is equal to RT-SOA. **b**, Performance of one participant. Saccade landing points are shown for each trial, arranged by PT (black dots, right y axis; data redrawn from ref. 39, experiment 1, participant 3). Targets were located at ±6°. The direction transition function (blue trace, left y axis) is a running histogram (bin size = 81 ms) of the proportion of correct saccades toward the stepped target as a function of PT. We computed it based on the shown saccade landing points. **c**, PT distributions in double-step (black trace) and distracter-step (pink trace) blocks performed by the same participant (data redrawn from ref. 39, experiment 1, participant 3). In distracter-step blocks, participants were instructed to always make a saccade to the first target, ignoring the step. **d**, Motor plans toward initial (black traces) and stepped (blue traces) target locations in 3 simulated double-step trials. A fast error (left), a correct saccade (middle), and a slow error or lapse (right) are shown. Shades indicate mean interruption interval. Triangles mark saccade onset. **e**, Simulated double-step responses (black dots; similar number of trials as in **b**) and direction transition function (blue trace; based on 20,000 trials). Correct and incorrect simulated saccades were assigned to +6° and −6° landing point values, respectively, with additional random scatter. **f**, PT distributions in double-step (black trace) and distracter-step (pink trace) simulated trials. Interruption and initial build-up rate parameters were the same for the two trial types. Gray shade shows the PT distribution for control, no-step simulated trials, which were not interrupted. For all data the SOA was constant at 120 ms.

Performance in the task is quantified via the direction transition function (Fig. 5b, blue trace). This curve describes the probability of making a correct saccade to the stepped target as a function of PT (Fig. 5a), which in this case is the interval between step onset and saccade onset (i.e., the amount of time available in any particular trial to view the target after it has stepped, where PT = RT - SOA). As can be seen for one example participant from the study by Buonocore and colleagues^39^ (Fig. 5b, blue trace), when the step is viewed for less than 100 ms or so, the resulting saccade is to the first, incorrect location, but as the PT increases beyond that mark, the likelihood of making a correct saccade to the stepped target rises steeply and then levels off. the transition from (fast) incorrect responses to correct ones occurs extremely rapidly, within 20 ms of PT or so.

Crucially, Buonocore and colleagues^39^ showed that the timing of the saccades in the double-step task is remarkably consistent with the target step producing robust saccadic inhibition of an initial motor plan toward the first target location. To see this, first consider the distribution of PTs (Fig. 5c, black trace), which includes both correct and incorrect responses (Fig. 5b, black points). The distribution is clearly bimodal, with a pronounced dip centered around PT ≈ 105 ms. This is, indeed, as if the target step had inhibited an ongoing saccade plan. To verify this interpretation quantitatively, Buonocore and colleagues^39^ did the following manipulation. In separate blocks of trials, they presented the same displays as in the double-step task but instructed the participants to ignore the step and make an eye movement to the initial target location (same sequence as in Fig. 5a, except with the correct saccade in the opposite direction). But note that this works as a classic saccadic inhibition paradigm, in which a salient distracter (the stepped target) must be ignored and, in addition, the actual target disappears. What they found was that the PT distribution from the distracter-step blocks (Fig. 5c, pink trace) was nearly identical to that in the double-step blocks; the saccades in both tasks produced the same unmistakable dip. This suggests that the interruption occurs in both cases and that it depends fundamentally on the stimulation conditions, regardless of how the abruptly-presented sensory information is used to guide the subsequent saccadic choice.

We performed computer simulations of these experiments (see Methods) to verify this conclusion, i.e., that the initial motor plan is interrupted in the same way in both conditions, except that, in one case (distracter-step trials) that first plan is meant to continue to threshold, whereas in the other (target-step trials) it is meant to be canceled and replaced by an alternate one. Motor plans toward the initial target location (Fig. 5d, black traces) were generated as in Fig. 1, by drawing from a Gaussian distribution a different build-up rate for each trial. These initial plans were always interrupted (i.e., the probability of interruption was equal to 1), unless they reached threshold before the onset of the pause, of course (Fig. 5d, left). The onset and offset of the interruption varied across trials, to produce smooth RT distributions. In the distracter-step blocks, after the interruption the initial motor plan simply continued to threshold. In contrast, in the target-step blocks a concurrent plan was launched at the beginning of the pause and the initial plan was canceled at the end of the pause (Fig. 5d, middle; this was true for correct responses; see below). The resulting PT distributions for the two conditions (Fig. 5f) had slightly different tails on the right side — a difference that was, in fact, consistent with the data of most participants in the study by Buonocore and colleagues^39^ (as with Fig. 5c). But more importantly, the two distributions displayed nearly identical dips. Although this may seem like a foregone conclusion, given that the initial plans and interruptions were statistically identical in the two cases, it is not. The simulations showed that, additionally, to produce such a tight match, the build-up rates of the initial and concurrent motor plans must have similar statistics.

The example participant made some errors at long PTs (Fig. 5b). These slower errors can be considered lapses, incorrect responses that cannot be attributed to insufficient viewing time. In the simulations, directional errors corresponding to lapses were generated by assuming that the cancelation of the initial plan is not 100% certain (Methods). That is, most often the initial plan is successfully interrupted and canceled, and a correct saccade to the stepped location is triggered (Fig. 5d, middle); however, sometimes that initial plan is interrupted but not canceled, resulting in an incorrect saccade (Fig. 5d, right). This mechanism produces lapses with the appropriate timing (Fig. 5e, lower group of dots with PT > 100 ms).

In this example, the same simulations reproduced not only the PT distributions but also the direction transition function (Fig. 5d, blue trace). The agreement depended both on dynamical features of the underlying motor competition process (e.g., the statistics of the build-up rates; see Methods) and on the parameters of the interruption. For example, making the interruption less reliable produced a less steep, more protracted direction transition function (data not shown). The main point exemplified by this task is that oculomotor performance can be strongly shaped by the interruption itself.

## Discussion

Because saccadic eye movements are planned continuously, the question arises of what to do when new visual information is detected. As a first approximation, there are three ways to proceed: (1) always let the ongoing plan continue (and deal with the new stimulus later), (2) always cancel the ongoing plan in favor of a new one congruent with the new stimulus, or (3) pause the ongoing plan, initiate a second, concurrent plan toward the stimulus, and deliberate to determine which one has a higher priority and should be executed next. Our review of the literature indicates that the latter, more flexible strategy is ubiquitous and robust, and our modeling results reveal two important aspects of it. First, the experimental manifestations of such strategy are accurately reproduced by a simple descriptive model in which the rise in oculomotor activity associated with a saccade plan is temporarily halted by a sudden-onset stimulus. Thus, although the quantitative details may vary across experimental conditions, the neuronal dynamics underlying such interruption are likely to be qualitatively the same. And second, the combined pause/concurrent-planning strategy is not necessarily optimal just because it is more flexible. For it to be advantageous, perception must be fast. Specifically, the advantage depends on how fast the new information can be interpreted relative to how fast the saccades can be programmed (equation (6)). When the deliberation is quick, on average it pays off to briefly pause the ongoing plan and initiate an alternative one right away *every time*. In this way, although ongoing plans to targets detected earlier are delayed, responses to novel, high-priority ones are rushed.

The results suggest a compromise whereby the RTs to high priority targets are shortened by a few tens of milliseconds per saccade while the responses to low priority targets are delayed by a comparable amount — that delay is the price to be paid for being ready to respond when novel information mandates an immediate action. The difference may seem modest, but could be critical for a cat that is patiently waiting for a mouse to spring out of its hiding place. In a competitive world the currency of survival is time itself, and mechanisms that enable slightly faster reactions may confer a vital advantage.

Besides providing a functional interpretation for the phenomenology associated with saccadic inhibition and attentional capture, the results are also important for models that aim to quantitatively link neuronal activity to oculomotor performance. As in the case of the double-step task, an interruption in motor planning of a few tens of milliseconds may introduce sizable behavioral and neurophysiological effects^6^, especially during urgent saccadic choices^5,16,58,59^.

### Deliberation and perceptual processing speed

Our calculations highlight the importance of sensory evaluation for saccade planning, and indicate that their speeds must match, i.e., the perceptual deliberation required to determine the priority of a novel stimulus must, on average, be shorter than the typical time between fixations (*Q* > *q*, equation (6)), which is about 200 ms in monkeys and humans^7,8,9^. Although many laboratory tasks create conditions in which sensory stimuli are judged for several hundreds of milliseconds or more (e.g., refs. 60, 61, 62), the perceptual decisions that characterize sensing in natural environments are likely to be much faster^63,64^. Specifically, the amount of time needed to make an accurate color discrimination is about 25–50 ms under urgent conditions, when oculomotor choices occur within a typical 200–250 ms RT window^5,16,20^. This number corresponds to the minimum amount of time that it takes for performance to transition from chance to 75% correct or above, so it is approximately equal to the deliberation time considered here, which excludes any transmission delays. Deliberation/processing times increase when stimulus discriminability goes down (Salinas and Stanford, unpublished results). But, on the other hand, the deliberation time is likely to be even shorter, closer to 20–30 ms, for a stop signal^65^, a single, salient stimulus that appears abruptly and is interpreted as a command to stop an action (e.g., a red traffic light). Thus, results based on urgent-choice paradigms reveal timescales for relatively easy, fast perceptual judgements of a few tens of milliseconds, consistent with the interruption durations inferred from saccadic inhibition experiments.

This timescale is also consistent with classic visual search experiments used to investigate attentional allocation^66,67^. In visual search tasks, the main measurement is the time needed to determine whether a target is present or absent in a display, and the key quantity characterizing the overall difficulty of the task is the search slope, i.e., the slope of the linear fit describing the RT as a function of the number of items in the display. For a given search task, the search slope corresponds directly to the deliberation time needed to identify each additional item as either target or distracter. Notably, search slopes are typically smaller than 10 ms/item when targets are highly salient or highly discriminable from non-targets, but even for difficult versions of these tasks, such as searching for a blue *H* among green *H*s and blue *A*s, deliberation times exceed 50 ms/item only rarely^66,67^ These numbers indicate that perception is normally quite fast — fast enough to satisfy the key constraint identified here, which makes the interruption/concurrent-programming strategy advantageous.

Still, given this constraint (*Q* > *q*), one might worry about the fact that intersaccadic intervals occasionally happen to be very short^8,68^ (*<* 100 ms), such as when a “corrective” movement is produced immediately after an incorrect saccadic choice^18,19,25,69,70^. In all likelihood, however, such short-latency responses are fast precisely as the result of concurrent motor programming. For instance, the latency of the correctives decreases as a function of PT in the double-step task^18,71,72^. This, together with evidence from single-neuron recordings in awake monkeys^19,72^, indicates that, during errors, the motor plan favoring the correct target keeps advancing up until the onset of the erroneous saccade (as in Fig. 5d, right), and likely contributes to the corrective movement that follows shortly thereafter. Thus, although extremely short intersaccadic intervals do occur, rather than negating the functional advantage of the interruption mechanism, they likely are a consequence of the concurrent motor programming that the interruption is proposed to enable in the first place.

### Temporal and spatial specificity

According to the proposed functional interpretation, the interruption should vary according to the spatial congruence between the ongoing motor plan and the abrupt-onset stimulus. When the unexpected stimulus appears close to or at the endpoint of the saccade that is currently being planned, there should be no interruption, as there is no competing motor plan. And indeed, the effect on RT has the expected spatial dependence. When a stimulus is precisely coincident with a saccade target, the eye movement is triggered sooner^55,56,68^, so the stimulus actually reinforces the ongoing motor plan. The decrease in RT goes away within a few degrees, as the location of the abrupt-onset stimulus deviates from the endpoint of the planned saccade vector, until the dip in the RT distribution emerges, and as the separation between the corresponding vectors approaches 180 °, it becomes most pronounced^47,56^. The interruption is manifest to the degree to which the ongoing and potential motor plans are in conflict with each other.

Congruence in the temporal domain is also important. Motor plans are reliably interrupted when a distant stimulus appears abruptly just before saccade onset, i.e., within the RT interval of the ongoing eye movement plan. When the time interval between the distracter and saccade onset is much longer (≥ 300 ms), as typically happens when a saccade target is presented 100 ms or more *after* the distracter, then the effect may go away or even reverse^56,73^. Our results are applicable specifically when the stimulus and the developing motor plan overlap in time, otherwise their interaction may be entirely different.

### Change of mind or change of plan?

The term “change of mind” has become popular for describing situations in which the neural activity predicting choice *A* is initially strong, but shortly thereafter the neural activity favoring option *B*, which was initially weak, gains momentum and prevails, so the choice is toward *B* ultimately^61,74,75^. These swings within single choice trials are sometimes explicitly described as “vacillations,” or signatures of indecision^74^. However, the current and previous results^5,6,16,58,59^ indicate that, rather than rare anomalies, these events must be extremely common, occurring as a natural and functionally useful aspect of the dynamics of motor circuits. The work of Thura and Cisek^76^ shows that, in premotor and motor cortex, motor plans favoring one or another arm movement are exquisitely sensitive to individual, quantized pieces of relevant sensory information, such that the prevalence of one plan over another can swing back and forth quite distinctly as pertinent evidence is presented over time. This is simply in keeping with the flexibility inherent to the definition of a “motor plan,” which implies that plans may be promoted, canceled, or otherwise changed as necessary up until the point of commitment^77^. That oculomotor circuits demonstrate a similar capacity to switch motor plans is perhaps less evident, insofar as saccades involve shorter timescales, fewer degrees of freedom, and stronger competition between them, making changes in saccade plans somewhat difficult to resolve (but see refs. 5, 6, 16, 78). Conceptually, however, they should not be surprising, as they simply reflect flexibility that is behaviorally advantageous. The transient interruption of ongoing motor plans is one specific gear in the oculomotor machinery that enables such flexibility.

## Methods

All simulations were performed using Matlab. In each simulated trial of the rise-to-threshold process, the firing rate variable *R* was integrated numerically by applying the update rule

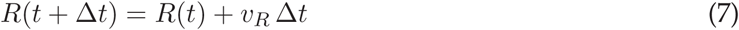

with a time step Δ*t* = 1 ms and a rate of change *v*_*R*_. The go signal was assumed to occur at *t* = 0, at which point *R* was equal to a baseline value *B* = 0. The saccade was assumed to be triggered when *R* reached a threshold Θ = 1000 arbitrary units (AU), at which point the RT was computed (the efferent delay was ignored, as it simply contributes a constant that can be consolidated with the afferent delay). Over the course of the trial, the rate of change took the following values

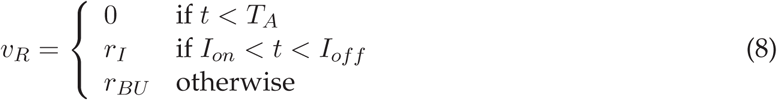

where *T*_*A*_ is the afferent delay, *I*_*on*_ and *I*_*off*_ are the onset and offset of the interruption, respectively, *r*_*I*_ is the build-up rate during the interruption, and *r*_*BU*_ is the nominal build-up rate of the process. For Figure 1, parameters were as follows. The afferent delay, *T*_*A*_, was drawn from a Gaussian distribution with a mean of 50 ms and a SD of 5 ms. The nominal build-up rate was drawn from a Gaussian distribution with a mean of 7.7 and SD of 1.9 AU/ms. The onset and offset times were drawn from independent Gaussian distributions with means of 192 and 228 ms, respectively, and equal SDs of either 0 (Fig. 1c, red trace) or 8 ms (Fig. 1d, red trace). For non-interrupted trials *r*_*I*_ = *r*_*BU*_, and for interrupted trials *r*_*I*_ = 0. The probability of interruption per trial, *p*_*I*_, was either 0 for simulations with no interruption (Fig. 1c, d, gray shade), 1 (Fig. 1c, red trace), or 0.7 (Fig. 1d, red trace).

For Figure 2, instead of integrating *R* numerically, equation (2) was applied directly. In each trial, first, a nominal RT without interruption, RT_0_, was sampled from the empirical RT distribution obtained with no distracter (Fig. 2b). For this, the published distribution^44^ was digitized and the numerical values were used as the input weights to the Matlab function randsample, which generates random samples from arbitrary distributions. Having drawn a sample RT_0_, the interruption was then computed for that trial. Interruption onset and offset times, *I*_*on*_ and *I*_*off*_, were drawn from a bivariate Gaussian distribution with a correlation of −0.8, means equal to SOA + 85 and SOA + 115 ms, respectively, and identical SDs of 14.3 ms. Depending on the probability *p*_*I*_, the interruption duration in the trial was either *q* = 0 or else *q* = *I*_*off*_ - *I*_*on*_. Finally, the simulated RT in the trial, RT_*I*_, was obtained by inserting RT_0_, *I*_*on*_, and *q* into equation (2).

Note that, for the results in Figure 2, the correlation parameter allowed us to consider a range of scenarios, going from cases in which the interruption duration stays relatively constant but its center point varies (correlation ≈ 1), to cases in which the center point remains constant but the duration of the interval varies (correlation ≈ −1). Together with *p*_*I*_ = 1, the parameter values just listed minimized the mean absolute difference between the empirical and simulated distributions, averaged across the 5 experimental conditions (for SOAs of 0, 20, 40, 60, and 80 ms).

For Figure 4, the RT associated with each individual rise-to-threshold process was calculated analytically using the interruption duration, *q*, the value of *α*, and the corresponding build-up rates (the initial one, based on *Q*, and the rate during the interruption, if nonzero). What was calculated numerically were the averages across trials as these parameters varied.

For Figure 5, equations (7) and (8) were used, and the parameter values for the initial motor plan and for the interruption were as follows: the afferent delay, *T*_*A*_, was drawn from a Gaussian distribution with a mean of 50 and SD of 10 ms; the initial build-up rate was drawn from a Gaussian distribution with a mean of 6.1 and SD of 1.7 AU/ms; the probability of interruption was *p*_*I*_ = 1; the build-up rate during the interruption was *r*_*I*_ = 0; the onset and offset times of the interruption were Gaussian samples with means of SOA + 53 and SOA + 157 ms, respectively, correlation equal to −0.8, and SD equal to 19.2 ms. The SOA was fixed at 120 ms, as was the case for participant 3 in the study by Buonocore and colleagues^39^. These values were identical between the simulated blocks of distracter-step and target-step trials. In distracter-step trials, the initial motor plan continued after the end of the pause, and the RT was recorded as the time when it reached threshold. In target-step trials, a concurrent plan was initiated during the pause. The build-up rate of this plan was drawn independently for each trial, with a mean of 6.2 and SD of 1.7 AU/ms. In correct distracter-step trials, the first plan was assumed to be canceled after the pause and the RT was recorded as the time at which the second, concurrent plan reached threshold (as shown in Fig. 5d, middle). In error trials, the cancelation failed, the first plan continued after the pause, and the RT was recorded as the time at which this plan reached threshold (as shown in Fig. 5d, right). Such failures occured when the build-up rate of the initial plan exceeded that of the concurrent plan by at least 1.5 AU/ms (which happened on 26% of the trials). Finally, whichever motor plan was still active after the pause, accelerated; that is, its build-up rate increased linearly between the end of the pause and threshold crossing. During distracter-step trials the initial plan accelerated at a rate of 0.03 AU/ms^2^, and during target-step trials the concurrent plan accelerated at a rate of 0.01 AU/ms^2^. The effect of these acceleration terms was to curtail the right tails of the RT distributions. They were not essential, as qualitatively similar results were obtained with zero acceleration; however, they are consistent with related saccadic choice models^5,16,58^, and slightly improved the fits between the experimental (Fig. 5b, c) and model data (Fig. 5e, f).

The Matlab code used to analyze and/or generate the data in the current study is available from the corresponding author on reasonable request.

## Acknowledgements

Research was supported by the NIH through grants R01EY021228 and R01EY025172 from the NEI.

## Notes

**Disclosures**: The authors declare no conflicts of interest, financial or otherwise.

